# Reconstitution of Arp2/3-Nucleated Actin Assembly with CP, V-1 and CARMIL

**DOI:** 10.1101/2024.05.13.593916

**Authors:** Olivia L. Mooren, Patrick McConnell, James D. DeBrecht, Anshuman Jaysingh, John A. Cooper

## Abstract

Actin polymerization is often associated with membrane proteins containing capping-protein-interacting (CPI) motifs, such as CARMIL, CD2AP, and WASHCAP/Fam21. CPI motifs bind directly to actin capping protein (CP), and this interaction weakens the binding of CP to barbed ends of actin filaments, lessening the ability of CP to functionally cap those ends. The protein V-1 / myotrophin binds to the F-actin binding site on CP and sterically blocks CP from binding barbed ends. CPI-motif proteins also weaken the binding between V-1 and CP, which decreases the inhibitory effects of V-1, thereby freeing CP to cap barbed ends. Here, we address the question of whether CPI-motif proteins on a surface analogous to a membrane lead to net activation or inhibition of actin assembly nucleated by Arp2/3 complex. Using reconstitution with purified components, we discovered that CARMIL at the surface promotes and enhances actin assembly, countering the inhibitory effects of V-1 and thus activating CP. The reconstitution involves the presence of an Arp2/3 activator on the surface, along with Arp2/3 complex, V-1, CP, profilin and actin monomers in solution, recreating key features of cell physiology.

## Introduction

Actin filaments polymerize at membranes, powering membrane movements and changes in cell shape as part of the process of cell migration ^1^. At plasma membranes, actin filament assembly powers outward force to create cell protrusions, and it powers inward force as part of endocytosis. For intracellular membrane vesicles and pathogens, actin filament assembly can power sorting and movement ^2–4^. Actin assembly dynamics can also promote vesicle fission or fusion, depending on the organelle ^5, 6^.

Cells regulate the location and timing of actin filament assembly with exquisite control ^7, 8^. First, cells suppress the spontaneous creation of filaments from subunits (i.e. nucleation) with actin monomer sequestering agents including profilin and thymosin beta-4 ^9, 10^. Second, cells contain regulated proteins, including Arp2/3 complex and formins, that promote the nucleation of actin filaments from subunits ^11, 12^. Arp2/3 complex is a major contributor to actin filament nucleation at the plasma membrane. Arp2/3 complex alone is essentially inactive, and membrane-bound activators, aka nucleation-promoting factors (NPFs), such as WASp and WAVE family proteins, help convert Arp2/3 complex to its active state. Arp2/3 complex binds to the side of an existing (mother) actin filament. This interaction also promotes Arp2/3 complex activation, and the activated molecule of Arp2/3 complex then binds actin monomers to create a new (daughter) filament, which can grow with a free barbed end at a 70-degree angle from the mother filament.

Biochemical reconstitution studies show that actin filament assembly and motility can be powered by a synthetic mixture of actin, Arp2/3 complex, capping protein (CP) and ADF/cofilin ^13–15^. In those studies, for each component of the mixture, an optimal activity level was observed; too much or too little of one component would lead to less than optimal actin assembly.

CP is intrinsically active for binding F-actin barbed ends under a wide variety of physiological buffer conditions. Inhibitors of CP bind directly to CP and inhibit its actin capping activity. The inhibitors include polyphosphoinositide lipids, the protein V- 1/myotrophin, and a set of proteins with CPI (CP-interacting) motifs ^16–18^. CPI-motif proteins are diverse; their sequences are completely unrelated to each other except for the presence of the conserved CPI motif ^16^. Among CPI-motif proteins, the CARMIL (capping protein, Arp2/3, myosin I linker) was discovered early ^19, 20^, and the CARMIL protein family has been studied rather extensively ^21–24^.

V-1 inhibits the actin capping activity of CP by sterically blocking the F-actin-binding site of CP ^25–28^. CARMIL and other CPI-motif proteins inhibit the actin capping activity of CP by an allosteric mechanism, binding at a distant site, away from the surface of CP that binds V-1 and F-actin ^28, 29^. CPI-motif proteins induce a conformational change in CP that propagates to the V-1 and actin-binding sites of CP, and this weakens the binding of CP to both V-1 and actin ^18, 30^.

V-1 and CP are present in cytoplasm at relatively high (microMolar) concentrations, and they bind with relatively tight affinity (nanoMolar values of K_D_) ^31^. V-1 and CP both appear to diffuse freely about the cytoplasm, and V-1 is in excess of CP ^31^. Thus, nearly all the diffusing CP in cell cytoplasm should be bound to and inhibited by V-1. CPI-motif proteins, including CARMILs, are present in far smaller quantities, and they are generally targeted to membranes via domains other than their CPI motifs ^18^. In a model put forth by Hammer and colleagues ^31^, a CPI-motif protein bound to a membrane at a certain location is proposed to activate CP locally by binding to CP / V-1 complex and promoting dissociation of V-1 from CP. Arp2/3 complex, activated by NPFs at the membrane location, can then nucleate actin polymerization in concert with active CP. This model is supported by the observation that a mutated form of CP, lacking the ability to bind a CPI motif, appears to lack function in cells, phenocopying a simple loss of CP ^32^.

We tested this hypothesis with a series of biochemical reconstitution experiments, a valuable approach in the field ^33^. Actin, profilin, V-1, and CP were added to beads coated with an Arp2/3 activator and the CPI motif from CARMIL. The beads thus mimic a location on the surface of the plasma membrane. The model predicts that V-1 will inhibit CP and thus prevent actin polymerization. The model also predicts that the presence of CARMIL will recruit CP to the bead surface, leading to activation of CP and thus actin polymerization at the surface. Our results demonstrate that all of these predictions are true.

## Results

### Capping protein is important for actin network formation and organization

Branched actin network assembly nucleated by Arp2/3 complex requires the presence of barbed-end capping activity ^33^, as demonstrated in early reconstitution studies ^14^. CP has provided barbed-end capping in many reconstitution studies ^33^. CP is important for actin comet tail formation from bacteria ^14^ and for breaking the symmetry of F-actin grown from a bead surface ^34, 35^.

To investigate the roles of the CP regulators V-1 and CARMIL, we first examined how the concentration of CP influenced the growth and overall organization of the F-actin network polymerized from beads. We coupled GST-VVCA, the potent Arp2/3-complex-activating domain of N-WASP ^36, 37^, to glutathione-conjugated polystyrene beads. The beads were incubated with actin (5 µM, 10% Alexa 488-labeled), profilin (5 µM), Arp2/3 complex (100 nM), and varying concentrations of CP. We observed the growth of the F-actin network by imaging the fluorescent actin, and we quantitated the area and total amount of fluorescence from the images (Figure 1, panels A-C). We observed a biphasic dependence of the growth of the actin network on CP concentration. In the absence of CP, a relatively small amount of F-actin accumulated in a symmetric ring surrounding the bead, and a larger cloud of low-intensity fluorescent F-actin formed symmetrically around the beads, extending a distance away from the bead. This symmetric cloud of low-density actin filaments is consistent with previous results ^35^. At low concentrations of CP (12.5 and 25 nM), the growth of F-actin from the beads was asymmetric, also consistent with previous observations ^35^. At 50 nM CP, the actin comet tail grew to a greater length. At higher concentrations of CP (200 nM and above), we observed less and less actin assembly, consistent with a role for CP of capping barbed ends and inhibiting net growth of actin filaments. The total actin fluorescence was also much lower at high CP concentrations (Figure 1, panel C), presumably reflecting a network with filaments of shorter length.

**Figure 1.**
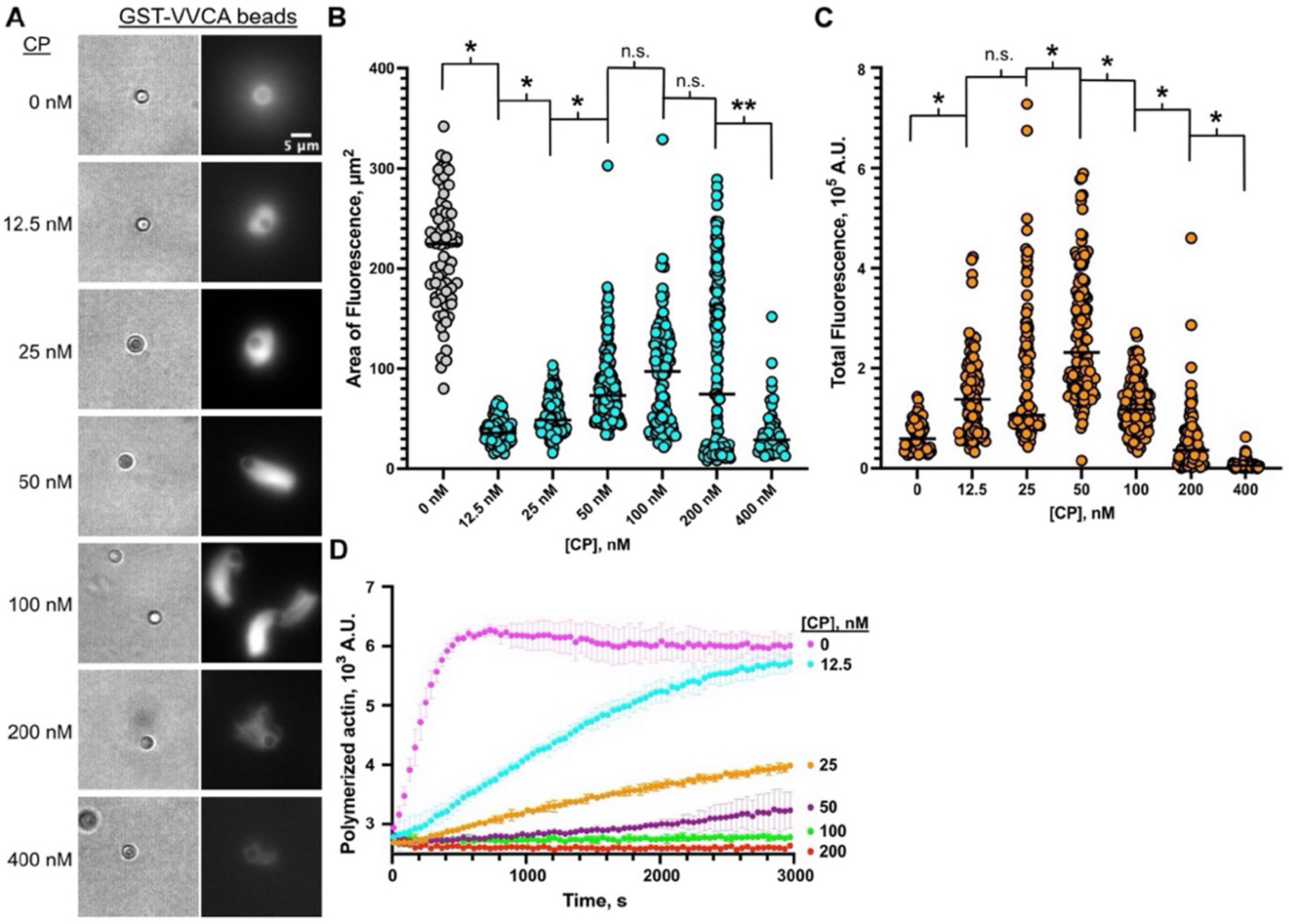
Actin network growth and organization is sensitive to CP concentration. **A.** Brightfield and fluorescence images of beads with actin networks formed with varying concentrations of CP. Beads coated with GST-VVCA were incubated with 100 nM Arp2/3 complex, 5 µM Alexa 488 actin, 5 µM profilin, and CP (concentrations as indicated). Images were collected at 30 min after the start of the reaction. **B.** The area of fluorescence around each bead was measured and plotted. Each dot is one bead. Grey dots represent beads surrounded by diffuse clouds of actin. Teal dots represent beads with asymmetric actin growth and tail formation. The presence of actin tails, the length of the tails, and the total area of fluorescence varied with CP concentration. The area of actin growth showed a biphasic dependence. Data from four independent experiments performed on different days are plotted. Non-parametric t-test (Mann-Whitney) p values with data compared to each CP concentration are indicated: * is p<0.0001, ** is p<0.01, and n.s. is not significant. **C.** Total fluorescence measurements of beads with varied CP concentration. Each dot is one bead. Results from four independent experiments performed on different days are plotted. Non-parametric t-test (Mann-Whitney) p values with data compared to each CP concentration are indicated: * is p<0.0001, n.s. is not significant. **D.** Pyrene-actin polymerization assays using beads demonstrate effective capping of actin in bulk as concentration of CP increases. This experiment was performed in triplicate; mean and standard deviation are plotted.

To complement the fluorescence imaging approach, we performed time-based pyrene-actin assays in solution in a plate-reader fluorometer, using the same components that were used for the imaging assays, including beads (Figure 1, panel D). In this setting, addition of CP decreased the amount of actin polymerization over time, and increasing CP concentrations caused further decreases in the time course of actin polymerization. This result supports the conclusion from the bead imaging experiments that high concentrations of CP limit the growth of actin filaments away from the bead. Considering all these results, we conclude that CP is important for modulating both the overall growth and the organization of branched filament networks.

### V-1 inhibits CP activity

V-1 sterically inhibits the actin capping activity of CP in biochemical assays, and the V-1 binding site on CP structurally overlaps with that of F-actin ^26–28^. In cell cytoplasm, the concentrations of V-1 and CP are high, their 1:1 binding affinity is tight, and the V-1 concentration exceeds that of CP, suggesting that nearly all cytoplasmic CP is bound to V-1 and thus inhibited for actin capping ^31^. We investigated how V-1 inhibition of CP alters actin network growth and organization by adding V-1 to bead-based reconstitution assays that contained an optimal level of CP (Figure 2).

**Figure 2.**
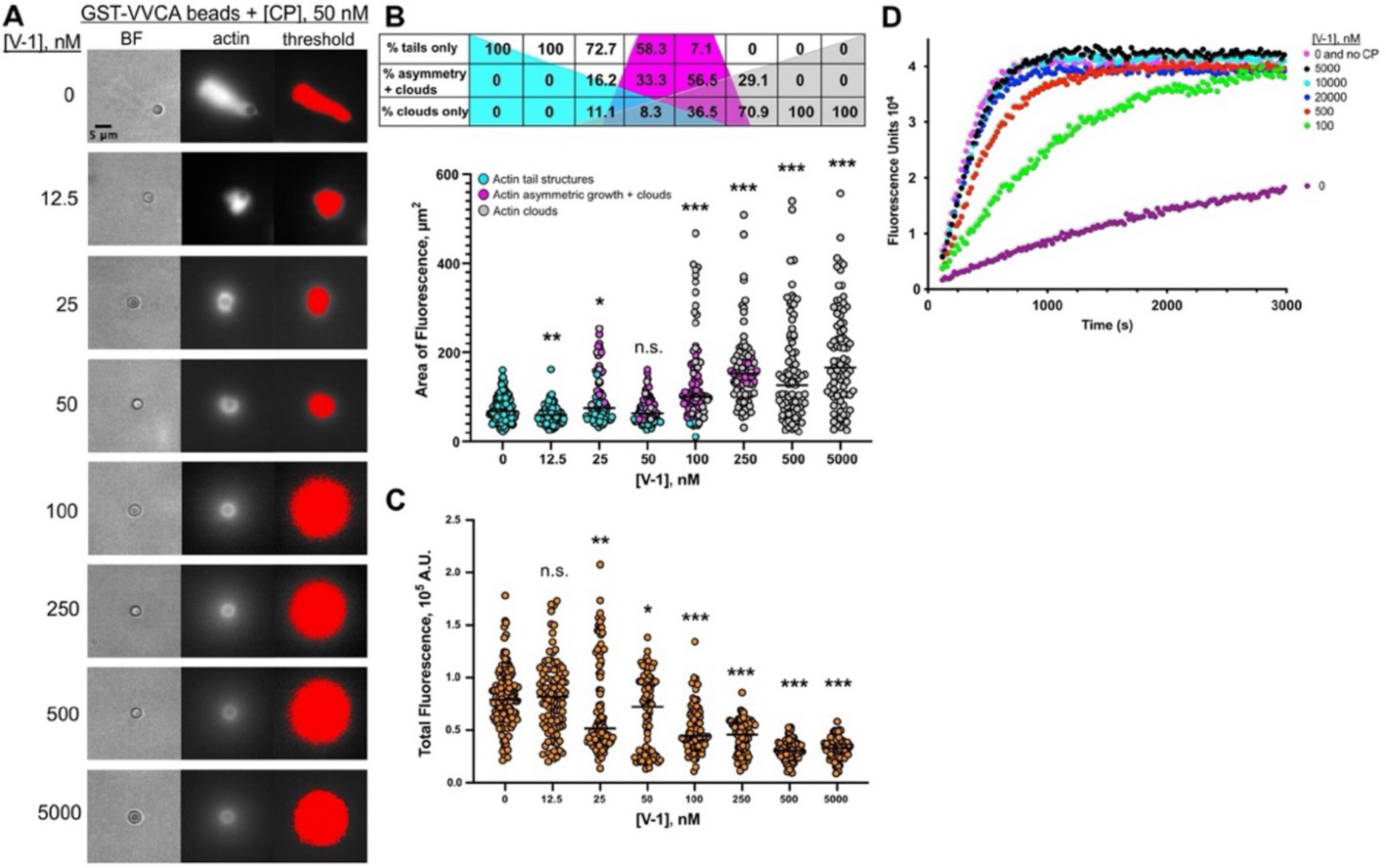
V-1 inhibits CP. **A.** Brightfield and fluorescence images of GST-VVCA-coated beads incubated with 100 nM Arp2/3 complex, 50 nM CP, 5 µM Alexa 488 actin, 5 µM profilin, and V-1 (concentrations as indicated). Threshold masks of the fluorescence images show low-intensity clouds of actin surrounding beads at higher V-1 concentrations. Images were collected at 30 min after reaction start. **B.** Area of actin fluorescence in these images is plotted, and percentages are listed. Teal-colored dots in the plot and teal shading in background of the table denote beads with actin tail structures. Pink dots and pink shading denote beads with asymmetric actin network growth and diffuse actin clouds. Grey dots and shading denote beads with symmetric growth of actin, appearing as a dense shell surrounding the bead surface, along with a diffuse cloud of actin. Data from three independent experiments are plotted. Non-parametric t-test (Mann-Whitney) p values with data compared to 0 nM V-1 conditions are indicated: * is p<0.02, ** is p<0.002, *** is p<0.0001, and n.s. is non-significant. **C.** Total fluorescence of the actin filament network is plotted, showing decreases with increasing concentrations of V-1. Data from three independent experiments is shown. Non-parametric t-test (Mann-Whitney) p values with data compared to 0 nM V-1 conditions are indicated: * is p<0.01, ** is p<0.001, *** is p<0.0001, and n.s. is non-significant. **D.** Pyrene-actin polymerization assays show that addition of V-1 inhibits the capping activity of CP in a dose-dependent manner.

Addition of low concentrations of V-1 shortened the actin tails (Figure 2, panels A and B), and higher concentrations of V-1 produced a transition from asymmetric tails to a mixture of asymmetric growth along with a diffuse cloud of fluorescence surrounding the beads. Further increases in V-1 concentration produced symmetric rings of intense actin fluorescence surrounding the beads, along with a diffuse cloud of fluorescence that extended farther away from the bead. The total area of the actin fluorescence (i.e., number of pixels with fluorescence intensity above background) increased with increasing V-1 concentration (Figure 2, panel B), and the summed total of actin fluorescence intensity over all pixels decreased with increasing V-1 concentration (Figure 2, panel C). This result indicates that the actin filament network is less densely organized in the presence of V-1. Together, these results – the area and total intensity of F-actin fluorescence, and the symmetry of the F-actin growth around the bead – are all consistent with V-1 decreasing the actin capping activity of CP.

We also examined the effect of V-1 on the overall growth of actin filaments by complementing the image-based assays with time-based pyrene-actin assays in solution using the same conditions, including the actin-nucleating beads (Figure 2, panel D). In the presence of 50 nM CP and the absence of V-1, F-actin growth was lower than growth in the absence of both CP and V-1 (purple curve vs fuschia curve), due to capping of barbed ends. With increasing concentrations of V-1, actin filament growth was restored (green to black curves), ultimately to the level observed with no CP and no V-1 (fuschia curve). These results indicate that CP decreases the growth of barbed ends, and that V-1 inhibits the actin capping activity of CP.

To extend our understanding of how V-1 inhibits CP, we used dual-color fluorescence imaging of CP and actin to examine the recruitment of CP to the bead and the resulting actin filament network. In the absence of V-1, CP was recruited to the bead surface, and the CP distribution mirrored that of the F-actin (Figure 3, panel A, top row). With the addition of a concentration of V-1 determined to be saturating in the pyrene-actin assays and the fluorescence bead imaging assays, we observed the loss of the recruitment of CP to the bead, as well as a corresponding shift from an asymmetric tail to a symmetric ring of F-actin with a diffuse cloud of F-actin surrounding the bead (Figure 3, panel A, bottom row).

**Figure 3.**
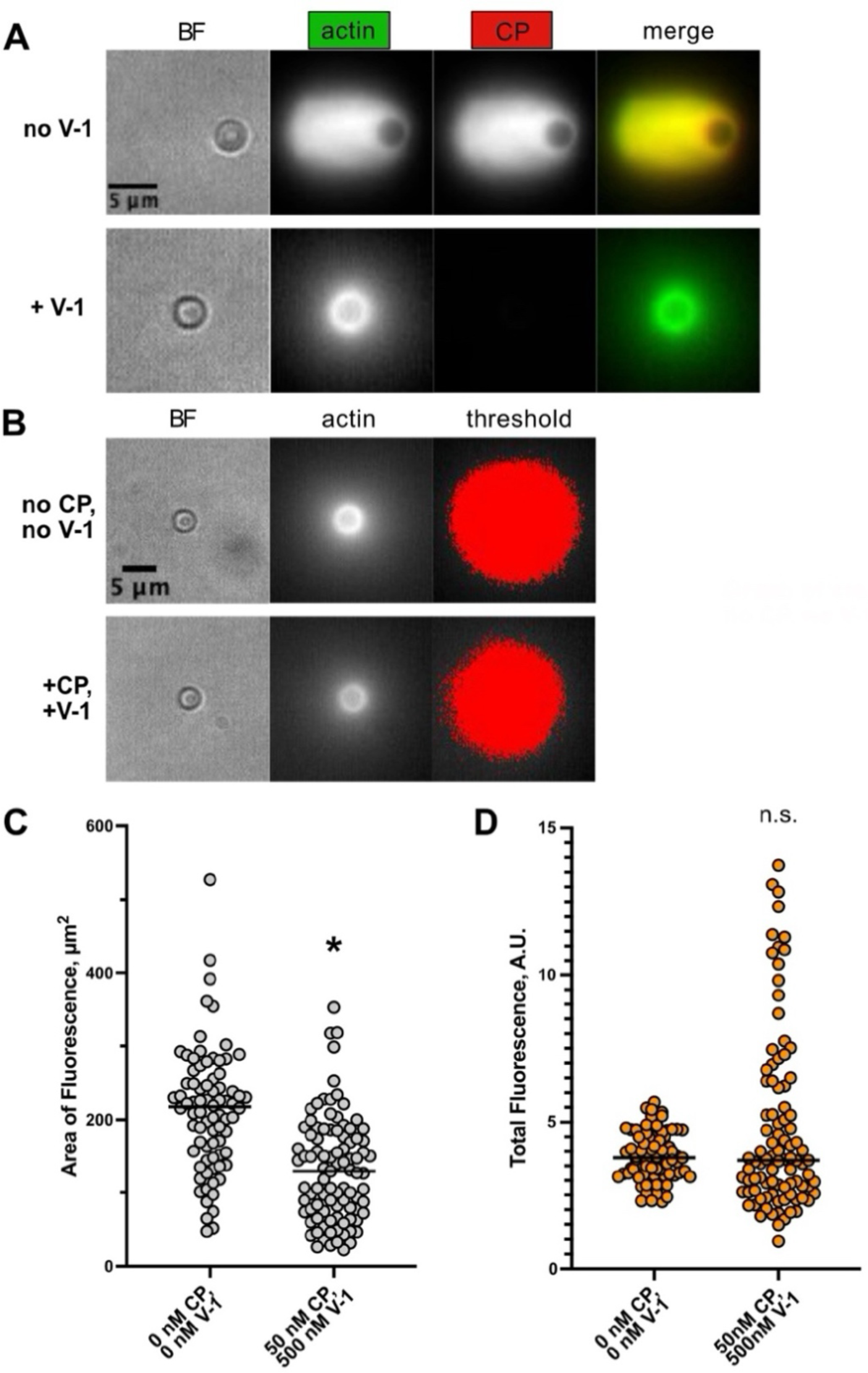
V-1 inhibits recruitment of CP to the actin filament network. Saturating concentration of V-1 caused the actin filament network to resemble that observed in the absence of CP**. A.** Brightfield (BF) and fluorescence images of beads GST-VVCA-coated beads mixed with 100 nM Arp2/3 complex, 50 nM Alexa 594 CP, 5 µM Alexa 488 actin, 5 µM profilin, and 500 nM V-1 (as indicated). **B.** Comparison of beads prepared in the absence of CP and V-1 with beads prepared with 50 nM CP and 500 nM (saturating) V-1. Threshold images show the area of the low-fluorescence-intensity actin clouds. **C.** Area of actin fluorescence for beads made with no CP and no V-1 compared with 50 nM CP and 500 nM V-1. Experiments were performed on three independent days, and all values are plotted. Non-parametric t-test (Mann-Whitney) p values are indicated: * is p<0.0001. **D.** Total actin fluorescence intensity measurements demonstrates that V-1 inhibition of CP results in about the same amount of polymerized actin as observed in the absence of CP. Experiments were performed on three independent days, and all values are plotted. Non-parametric t-test (Mann-Whitney) p values are indicated: n.s. is non-significant.

The actin filament organization in the presence of saturating V-1 conditions resembled that observed when no CP was included in the reaction mixture (Figure 3, panel B). We quantitated the area of actin fluorescence and the total actin fluorescence intensity under both conditions. The area of the actin fluorescence surrounding beads in saturating V-1 conditions was less than that in conditions where no CP was included in the reaction (Figure 3, panel C). We speculate that this result is due to a small amount of CP that is not bound to V-1 at any given time, providing for a low level of CP activity. The total actin fluorescence, reflecting the summed amount of F-actin polymer mass, was the same for both conditions (Figure 3, panel D). Together, these results demonstrate that V-1 acts as an inhibitor of the actin capping activity of CP in this setting.

### Surface-bound CARMIL activates CP at the bead

CARMILs are CPI-motif-containing proteins able to inhibit the barbed-end actin capping activity of CP and to promote dissociation of CP from capped barbed ends, i.e. “uncapping.” ^23, 38–40^. V-1 inhibits CP by sterically blocking the actin binding site of CP ^26–28^. In contrast, CPI-motif proteins bind to the opposite side of CP and allosterically induce a conformational change in the surface that binds F-actin and V-1 ^16, 30, 41^. The two effects of this conformational change are to increase the dissociation rate for V-1 from CP / V-1 complex and to weaken the binding affinity of CP for the barbed end of the actin filament ^31, 42^.

Initial studies showed that the biochemical effects of CPI-motif proteins on CP are to decrease its actin capping activity and promote uncapping ^23, 40^. However, CP in complex with a CPI motif was found to be able to bind to actin barbed ends, albeit with a weaker binding affinity than that of CP alone ^23, 42^. Together, these biochemical observations led to the proposal by Hammer and colleagues that CP in cell cytoplasm is bound to V-1, and thereby sequestered in an inactive state until the CP / V-1 complex encounters a CPI-motif protein, such as CARMIL ^31^. The effect of CPI-motif binding is to promote release of free CP from V-1, allowing CP to associate with barbed ends nucleated by Arp2/3 complex near a membrane. We tested this model in our bead-based reconstitution system.

To mimic the membrane localization of CARMIL, we used a GST fusion of a segment of CARMIL1, “CBR115” in our nomenclature, that binds CP tightly and has strong biochemical effects on CP actin capping activity ^39^. GST-CBR115 was coupled to glutathione-functionalized beads along with a GST fusion of VVCA, the Arp2/3-activating fragment of N-WASP. As presented above, when neither V-1 nor CARMIL1 were present, the actin filament network grew as asymmetric tails, with CP at the bead surface in a distribution matching the actin fluorescence (Figure 4 panel A, top row). With saturating amounts of V-1, we observed symmetric actin growth around the bead with a diffuse cloud of actin, and little to no CP was recruited either to the bead or to the F-actin network (Figure 4, panel A, middle row). However, with addition of the CBR-115 CARMIL fragment at the bead surface, the resulting actin network showed asymmetric growth from the bead surface, and CP was once again localized to the bead and to the actin network (Figure 4, panel A, bottom row).

**Figure 4.**
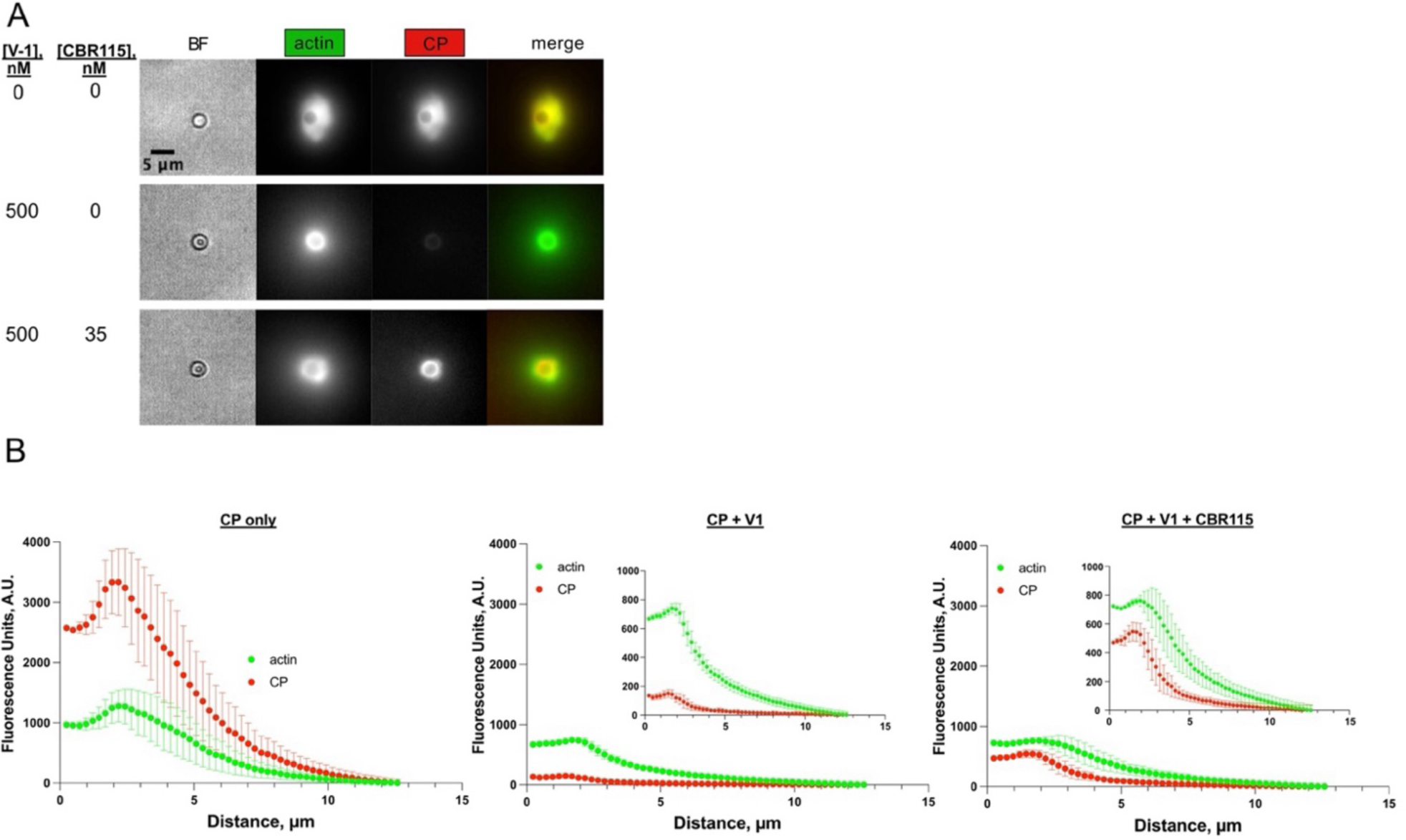
CARMIL on the bead surface recruits CP to the surface and influences actin network growth and organization. **A.** Beads coated with either GST-VVCA or GST-VVCA plus GST-CBR115 were mixed with 100 nM Arp2/3 complex, 50 nM Alexa 594 CP, 5 µM Alexa 488 actin, 5 µM profilin, and 500 nM V-1 (as indicated). **B.** Radial scans of fluorescence intensity for beads shown in panel A. Zero distance is the bead center. Actin in green, CP in red. The values for fluorescence on the y-axis are directly comparable among the three panels because the images were collected in one experiment with identical microscope settings. The inset plots are on a smaller y-axis scale to illustrate the differences in intensity and variance among the samples.

To quantitatively analyze the growth of the F-actin network and the recruitment of CP, we performed radial scans of the intensities of the F-actin and CP fluorescence images. We averaged pixel intensity values as a function of distance from the bead center for all directions (Figure 4, panel B). Radial scans in the sample with CP alone, with neither V-1 nor CARMIL, showed similar fluorescence distributions and peaks for actin and CP (Figure 4, panel B left side).

With saturating amounts of V-1, radial scans showed that the level of actin fluorescence was lower than that observed in the absence of V-1 (Figure 4, panel B middle, and noted above for Figure 2). The level of CP decreased nearly to zero, as well. In addition, the radial scan traces for both actin and CP contained much lower variance in the presence of saturating V-1 compared to no V-1 (Figure 4, panel B left vs. middle). We calculated a metric for the radial scan traces of the actin by taking the sum of the standard deviations divided by the sum of the intensity values, which reflects the asymmetry of the distribution around the bead and, by extension, the level of active CP. We refer to this metric as the “variance metric” here. Bead samples containing V-1 had a median variance metric value of 0.0134 ± 0.004 (Suppl. Figure 1).

Radial scans of beads containing the CBR-115 CARMIL fragment displayed a higher level of variance for both actin and CP than for the V-1 alone conditions, reflecting the asymmetric growth of the F-actin network near the bead surface (Figure 4, panel B right side). The variance metric value from the radial scans of the actin for 35 nM CBR115 CARMIL bead samples was 0.0178 ± 0.011 (Suppl. Figure 1). The CP localization did not mirror exactly that of the F-actin, as observed in the absence of V-1 and CARMIL, but it did extend away from the bead surface. In our interpretation, the CP distribution reflects the concentration of CP near the bead surface due to recruitment by CARMIL. CP did not interact with all barbed ends of the F-actin network, due to the presence of V-1, but it remained attached to barbed ends after dissociation from CARMIL.

Another approach we used to quantify radial variance and objectively assess the level of asymmetric actin growth from the bead surface was to measure the spread of distances from the center to the perimeter of an object, as defined by thresholding. If the radial variance is low, the distances vary less, and the shape is smoother and more symmetric. A higher radial variance reflects variation in distances from the center, which can result from the object having an irregular shape or rough surface. We found that samples containing V-1 with CBR115 CARMIL had a higher radial variance than samples containing V-1 without CBR115 CARMIL (Suppl. Figure 2).

From these results, we conclude that CARMIL recruits or targets CP to the bead surface by binding to soluble CP / V-1 complex and promoting dissociation of V-1 to activate CP for capping actin. Activated CP, either bound to CARMIL at the surface or released into solution, is then able to interact with actin nucleated by N-WASp (VVCA) and Arp2/3 complex, thereby promoting F-actin growth and influencing the organization of the filament network.

To extend our analysis, we examined how the effects of CARMIL (GST-CBR115) varied with the amount of GST-CBR115 bound to beads (Figure 5). First, we validated that incubating beads with increasing amounts of GST-CBR115 led to increased amounts of GST-CBR115 attached to the beads, using Western blot analysis to measure the amount of bound GST-CBR115 per bead (Suppl. Figure 3). As expected, the amount of GST-CBR115 on beads increased with the concentration of GST-CBR115 added to the reaction, varying from zero to 21,000 µm^-2^ (Suppl. Figure 3).

**Figure 5.**
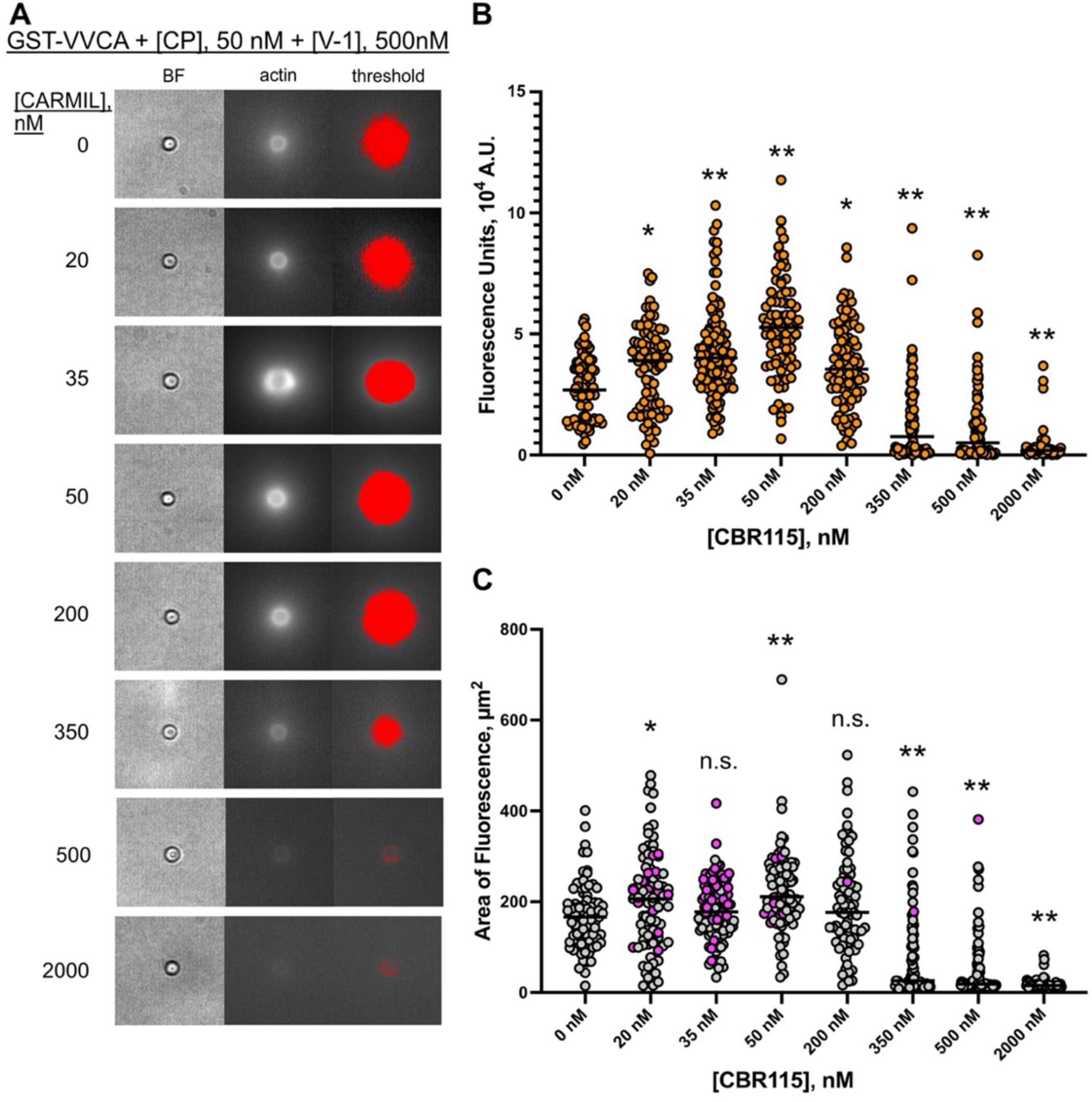
Ratio of CARMIL to CP determines the level of F-actin growth and overall density of actin network. **A.** Images of beads coated with GST-VVCA (N-WASP) and GST-CBR115 (CARMIL) at indicated concentrations and mixed with 100 nM Arp2/3 complex, 50 nM CP, 5 µM Alexa 488 actin, 5 µM profilin, and 500 nM V-1. Images collected at 30 min. **B.** Total fluorescence intensity of the actin filament network surrounding each bead. Results from three independent experiments were combined and are shown here. Non-parametric t-test (Mann-Whitney) p values with data compared to 0 nM CBR115 conditions are indicated: * is p<0.001, and ** is p<0.0001.**C.** Area of fluorescence around each bead was determined following thresholding of the image. Pink dots indicate beads that have asymmetric actin network growth around the bead and diffuse actin clouds. Grey dots indicate beads with diffuse actin clouds surrounding a symmetric shell of actin around the bead. Results from three independent experiments were combined and are shown here. Non-parametric t-test (Mann-Whitney) p values with data compared to 0 nM CBR115 conditions are indicated: * is p<0.01, ** is p<0.0001, n.s. is non-significant.

We measured the amount of N-WASP (GST-VVCA) on the bead surface to ensure that the amount of nucleation promoting factor (NPF) remained constant while the amount of attached GST-CBR115 increased. Plain GST was added to each reaction mixture in amounts designed to maintain a constant concentration of total GST (plain GST plus GST fusions) in solution in the bead coating process, as detailed in Methods. The density of GST-VVCA was found to be relatively constant at ∼1700 molecules per µm^2^ of bead surface (Suppl. Figure 4) while the GST-CBR115 density varied.

Small amounts of CBR115 CARMIL were sufficient to produce notable effects on the organization of the actin filament network around the bead (Figure 5, panel A). In addition, those relatively small concentrations of CBR115 CARMIL produced an optimal amount of actin growth as determined by total actin fluorescence (Figure 5, panel B). However, the total area of actin fluorescence around the bead increased only slightly, albeit to a statistically significant degree (Figure 5, panel C). When we analyzed only the area of actin fluorescence that was brightest around the bead, excluding the actin cloud, we observed an increase in actin area with the concentrations of CBR115 CARMIL (35 and 50 nM) that displayed the highest actin fluorescence (Figure 5, panel B and Suppl. Figure 5). The dependence of total actin fluorescence on CBR115 CARMIL was biphasic, with higher amounts of CBR115 CARMIL leading to lower total actin fluorescence and smaller areas of actin fluorescence (Figure 5, panels A-C). This biphasic dependence is similar to the results above with titration of CP (Figure 1), where higher levels of CP greatly diminished growth of the actin filament network growth.

As noted above with respect to Figure 4, the shape of the brightest actin fluorescence surrounding the beads became asymmetric when CBR115 CARMIL was present on the beads, with the peak level of asymmetry observed for 35 nM CBR115 (Figure 5, panel C, magenta-colored points). We analyzed the shape of the most intense fluorescence closest to the bead surface by calculating roundness as a metric for symmetry/asymmetry. The value of roundness for a circle is 1 (Suppl. Figure 6, panel A). Beads with no CBR115 had a roundness value of 0.96 ± 0.02. Beads with CBR115 had lower roundness values, with 35 nM CBR115 displaying the largest distribution. In contrast, when we analyzed actin fluorescence including the less dense actin clouds, CBR115 caused only a slight decrease in roundness (Suppl. Figure 6, panel B). Thus, CBR115 CARMIL influences the actin network closest to the bead surface, with less effect on presumably uncapped filaments that grow away from the bead surface and into the sparse actin cloud. This interpretation is consistent with CBR115 CARMIL recruiting and activating CP at the bead surface.

To test this conclusion further, we examined recruitment of CP to the beads with increasing CBR115 CARMIL, by measuring the fluorescence of Alexa 594-CP on the bead (Suppl. Figure 7, panels A & C). As the concentration of CBR115 increased, the amount of recruited CP increased (Suppl. Figure 7, panel C). In addition, we confirmed the increase in CBR115 CARMIL on the bead surface using immunofluorescence with antibodies to CBR115 (Suppl. Figure 7, panel D). Overall, we conclude that surface-bound CARMIL recruits CP and regulates its activity at the bead surface, resulting in altered actin organization and growth.

## Discussion

The most important conclusions from this study are 1) a CPI-motif protein (CARMIL) can recruit CP on a surface, 2) the recruited CP is active for actin capping because the allosteric effect of CARMIL promotes the dissociation of the steric inhibitor V-1, and 3) the combined effect is to promote Arp2/3-nucleated actin polymerization at the surface. The key finding is that the allosteric effect of CARMIL on CP counteracts the steric blocking inhibitory effect of V-1. The results support the model diagrammed in Figure 6. In this model, first proposed by Hammer and colleagues ^31^, the activity of CP is inhibited by the presence of stoichiometric amounts of the protein V-1 in solution. CARMIL tethered at the surface of a bead, which mimics a cell membrane, recruits CP to the bead surface and promotes dissociation of V-1, thus activating CP near actin filament barbed ends. Capping activity promotes Arp2/3-based nucleation because barbed ends can directly interact with WH2 domains, which are present in N-WASP. If the barbed ends are capped, the interaction with N-WASP is blocked, promoting nucleation by Arp2/3 complex.

**Figure 6.**
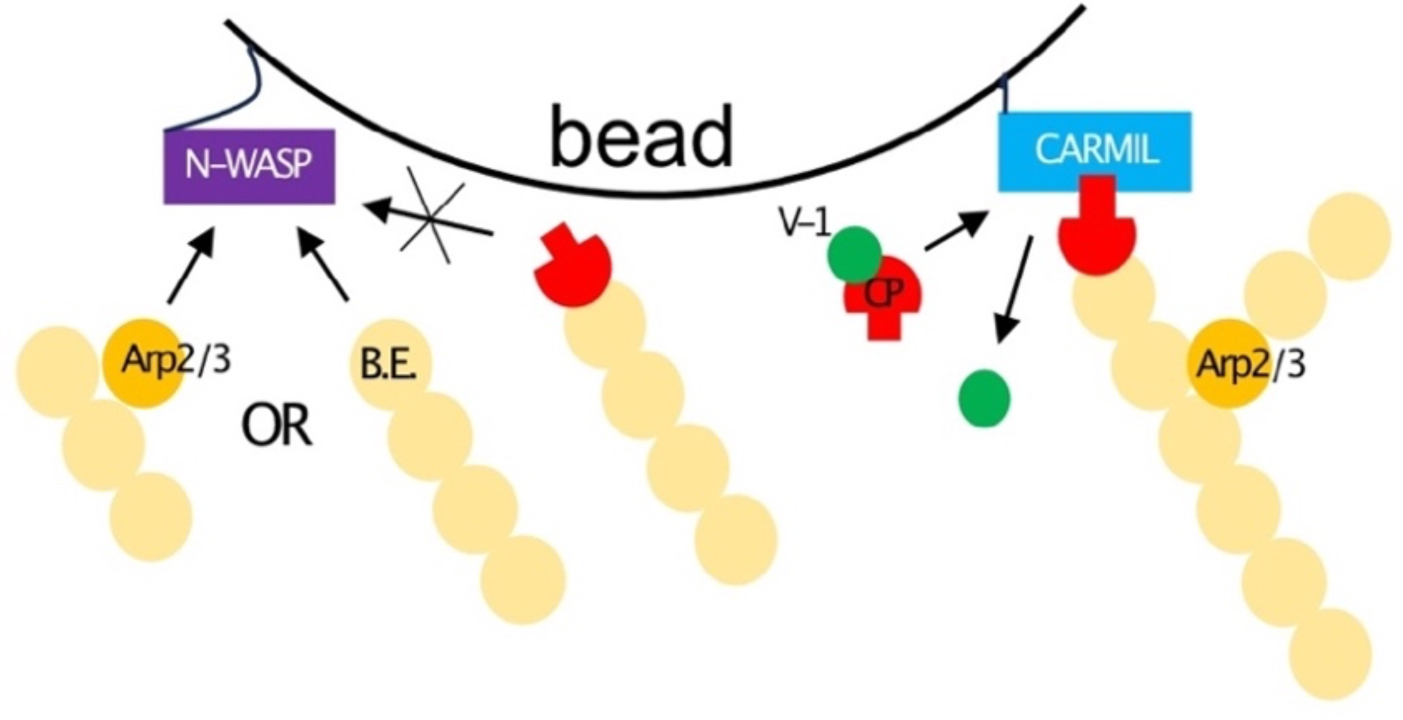
Model. The bead surface mimics the cytoplasmic face of a cell membrane. The surface displays an active fragment of the nucleation promoting factor (NPF) N-WASP (VVCA) along with an active fragment of the CPI-motif protein CARMIL1. VVCA recruits Arp2/3 complex from solution and promotes its activation for nucleation of actin polymerization. Free actin filament barbed ends (B.E.) are also able to interact with the VVCA domain of N-WASP, competing with Arp2/3 complex. CARMIL1 interacts with CP and recruits inactive CP / V-1 complex from solution. The effect of CARMIL1 is to promote the dissociation of V-1, which uncovers the actin capping surface of CP. Active CP promotes the ability of Arp2/3 complex to nucleate and assemble branched networks of actin filaments.

### Role of CP in Regulation of Actin Polymerization

The creation of free barbed ends of actin filaments by nucleation from actin subunits is a major requirement for the growth of actin filaments in cells ^1, 11, 12^. Cells use several regulated mechanisms to promote nucleation and create free barbed ends at specific times and locations. These free barbed ends elongate as actin filaments, and the assembly of those filaments powers and directs changes in cell shape and movement. Major actin nucleating proteins in cells include Arp2/3 complex and the formin family ^43, 44^. These nucleator proteins are recruited and regulated spatially and temporally by endogenous cellular proteins ^45–47^ and by proteins derived from pathogenic organisms ^48, 49^.

CP is important in cells for actin assembly and for actin-based motility ^8, 12, 17^. Biochemically, CP caps the barbed ends of actin filaments. The capping of free barbed ends of actin filaments by CP in biochemical assays with purified proteins in vitro suggested that CP might inhibit actin assembly in cells ^50^. Indeed, that conclusion was supported by early work altering the levels of CP in cells ^51^. However, biochemical reconstitution experiments with purified proteins subsequently revealed that low concentrations of CP promote, not inhibit, actin polymerization from barbed ends that are nucleated by Arp2/3 complex ^14, 35^. In these reconstitution studies, the dependence of actin polymerization on CP concentration exhibited a bell-shaped curve. There was no assembly in the absence of CP, actin assembly increased with increasing CP, peaking at a maximum level at an optimal CP concentration. With increasingly higher concentrations of CP, actin assembly decreased, ultimately to low levels ^14^. We likewise observed these trends in our biochemical reconstitution assays, showing the importance of having a balanced ratio of active capping protein to promote Arp2/3-based actin network growth.

### Combined Roles of V-1 and CARMIL in Regulation of CP

In cell cytoplasm, CP is present at relatively high (microMolar) total concentrations ^31, 50^, and CP binds to barbed ends with high affinity (nanoMolar K_D_) ^52^. These observations suggested that a newly created free barbed end in cells might add actin subunits for a period of time, and become capped over time as diffusing CP encounters and bind to the barbed end. The simplicity of this model was called into question by the discovery of two major classes of CP regulators, both of which bind directly to CP and lessen the ability of CP to bind barbed ends ^17^.

One major regulator of the actin capping activity of CP in cells is the protein myotrophin / V-1 ^26^. In biochemical assays with purified proteins, V-1 and CP bind with high affinity at 1:1 stoichiometry, and the dissociation rate for the V-1 / CP complex is slow ^26^. The binding site for V-1 on CP sterically overlaps with that of the barbed end, accounting for the competitive inhibition of CP’s actin capping activity by V-1 ^27, 53^. Cell cytoplasm has a high concentration of V-1, comparable to or greater than that of CP; thus, nearly all cytoplasmic CP should be sequestered as inactive by the action of V-1 ^31^.

Another major mode of regulation of CP is provided by the direct 1:1 binding of a set of proteins with “CP-interacting (CPI)” motifs ^16^. CPI motifs are found in several families of proteins that are otherwise completely unrelated ^18^. Biochemically, CPI-motif proteins bind to CP at a site distinct and away from the actin binding site. The effect of CPI-motif binding is to allosterically decrease the ability of CP to bind and thus cap actin barbed ends; the allosteric effect is mediated by conformational changes on the actin-binding surface of CP ^30, 41, 42^.

These conformational changes are modest, and the actin capping ability of CP bound to a CPI motif is lessened by a substantial amount but not abolished ^23, 42^. This decrease in capping activity may be physiologically relevant. The dynamics and level of capping activity provided by a CPI-motif / CP complex may be more suited for control of barbed ends in living motile cells than free CP alone.

When CPI-motif proteins bind to CP and induce a conformational change in the actin-binding site of CP, the conformational change also affects the interaction of V-1 with its overlapping binding site ^31, 42^. CPI-motif proteins bind CP / V-1 complex, and this interaction increases the rate of dissociation of V-1 from CP ^31, 42^. Together, these biochemical properties suggest that a CPI-motif protein in cells might convert CP from its inactive V-1-bound form to an active form, which is able to cap barbed ends, albeit with activity less than that of free CP ^31^.

This model was proposed by Hammer and colleagues, based on their observations of biochemical properties CP and V-1, along with measurements of cellular concentrations of CP and V-1 ^31^. CPI-motif proteins, including the archetypal CARMIL family, are generally bound to cellular membranes, including the plasma membrane; thus, this model proposes that membrane-bound CARMIL causes local activation of CP, which promotes actin polymerization from free barbed ends that were created by the nucleation activity of Arp2/3 complex. Arp2/3 complex also requires activation by membrane-bound proteins, such as the WASp and WAVE protein families ^46^. Thus, actin assembly at membranes involves two independent promoting mechanisms.

Here, we tested the model using a biochemical reconstitution system. To mimic the plasma membrane, we employed beads whose surface displayed an active fragment of an Arp2/3 nucleation-promoting factor and an active fragment of CARMIL. Both fragments were prepared as GST fusion proteins and thus bound to the surface of the glutathione-coupled bead in a specific manner. With the beads in solution, we added purified Arp2/3 complex, CP, and V-1. Actin monomers were then added, initiating the polymerization of actin filaments from barbed ends created by the nucleation activity of Arp2/3 complex. Profilin was also included as a physiological cellular component that suppresses spontaneous nucleation of actin filaments from monomers.

Together, our results support the key features of the Hammer model. First, bead-bound CARMIL recruited CP to the bead surface. Second, addition of V-1 inhibited the ability of CP to promote Arp2/3-based actin polymerization from the bead. Most important, the presence of CARMIL on the bead surface was able to overcome the inhibitory effect of V-1, allowing actin polymerization to proceed with nucleation by Arp2/3 complex and promotion by active CP.

Among our results, we note that growth of the F-actin network under conditions where CP was liberated from V-1 by CARMIL was not as robust as when CP alone was included without V-1 or CARMIL (Figure 4). As mentioned above, biochemical studies have shown that CARMIL-bound CP still retains some barbed-end capping activity to a limited extent. This level of capping activity might be expected to slow the growth of actin filaments (Figure 6). Additionally, as we titrated higher amounts of CBR115 CARMIL on the beads, we further suppressed actin filament growth, presumably due to activating more CP at the bead surface and inhibiting growth of barbed ends at the bead.

### Arp2/3 Complex and CP Regulation of Actin Barbed Ends

Previous studies have demonstrated the importance of CP in Arp2/3-based nucleation ^35, 54^. In particular, actin barbed ends can bind directly to the WH2 domains of Arp2/3 nucleation promoting factors, thus decreasing the activation of Arp2/3 complex and the amount of nucleation ^54^. CP activity can increase Arp2/3-based nucleation because capping of barbed ends prevents those ends from interacting with a WH2 domain. In the case of our studies, two WH2 domains are found in the GST-VVCA N-WASP fragment used to bind and activate Arp2/3 complex. Therefore, VVCA can bind to and activate Arp2/3 complex, or it can bind to barbed ends of filaments (Figure 6). When CP is bound by V-1, that CP cannot bind to barbed ends and thus cannot counteract the interaction of WH2 with barbed ends.

We observed a decrease in the amount of F-actin on beads incubated with saturating amounts of V-1, as well as a change in the organization of the F-actin network. This result may be due to a decrease in nucleation because more free barbed ends are available to compete for WH2 binding. Ultimately, one might envision a feedback mechanism in which filament ends can stay capped, but filaments grow slowly if CP dissociates from the filament end (perhaps due to its interaction with CARMIL), with cycles of capping and uncapping of barbed ends causing cycles of promoting and competing for nucleation by Arp2/3 complex. In this way, the growth and organization of the actin network can be fine-tuned by cells with CPI-motif proteins.

### Open Questions and Future Directions

Several questions about the physiological functions of CPI-motif proteins remain open. Our results here suggest avenues for addressing those questions. First, one issue is the evolutionary specialization of the biochemical properties of the conserved CPI motifs found in different families of CPI-motif proteins ^30, 42^. A related issue is the role of the CSI motif found in tandem with the CPI motif in the CARMIL family but not other CPI-motif proteins ^16^. The presence of the CSI motif makes CARMILs bind and inhibit CP better than other CPI-motif proteins ^29, 39^. CARMILs are also distinct from other CPI-motif proteins in having a membrane-binding domain in tandem with its CPI and CSI motifs ^55^. The presence of the membrane-binding domain increases the CP-binding activity of the CPI and CSI motifs in studies of proteins in solution ^29, 39^, but the effects in the presence of a membrane are not known.

Another important question is whether (and if so, when) CPI-motif proteins promote uncapping of barbed ends that are capped by CP in cells. In biochemical assays, this activity is prominent, especially for CARMIL family proteins due to their CPI, CSI and membrane-binding domains ^38^. The physiological relevance of this uncapping activity may be to promote the turnover of actin filaments found in stable arrays, such as sarcomeres ^56–58^ and stereocilia ^59, 60^.

## Conclusions

CARMIL can activate CP at membranes by dissociating CP from V-1, thereby promoting actin polymerization nucleated by Arp2/3 complex. In the absence of V-1 and CARMIL, low levels of active CP lead to polarized and structured growth of the actin network, but too little or too much CP leads to unstructured networks or overall restriction of actin growth. These effects of CP are most likely due to changes in Arp2/3 complex-based nucleation activity because of competition of barbed ends with WH2 domains. CP can bind actin when CARMIL is present, with a weaker association to the filament barbed end relative to CP alone. The system of interactions provides several points of feedback control.

## Conflict of interests

The authors declare no conflict of interests.

## Acknowledgments

We are grateful to members of our laboratories for advice and assistance. This research was supported by NIH grants R35 GM118171 and R35 GM144082 to J.A.C.

## Supplementary Figures

**Supplementary Figure 1.**
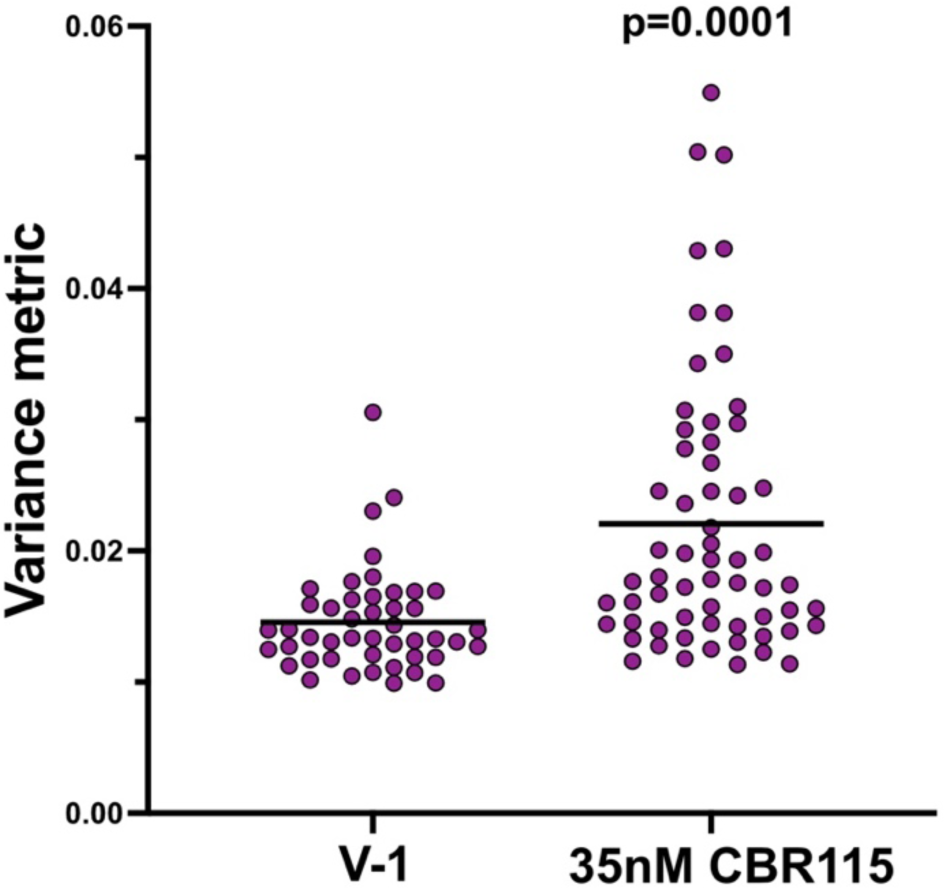
Variance increases with CARMIL indicating asymmetric actin growth around the bead. V-1 alone is compared with CBR115 CARMIL plus V-1. The variance metric for actin fluorescence was calculated for two independent experiments, showing that CBR115 CARMIL plus V-1 increases the amount of asymmetric actin growth around the bead compared with V-1 alone. Non-parametric t-test (Mann-Whitney) p value shown.

**Supplementary Figure 2.**
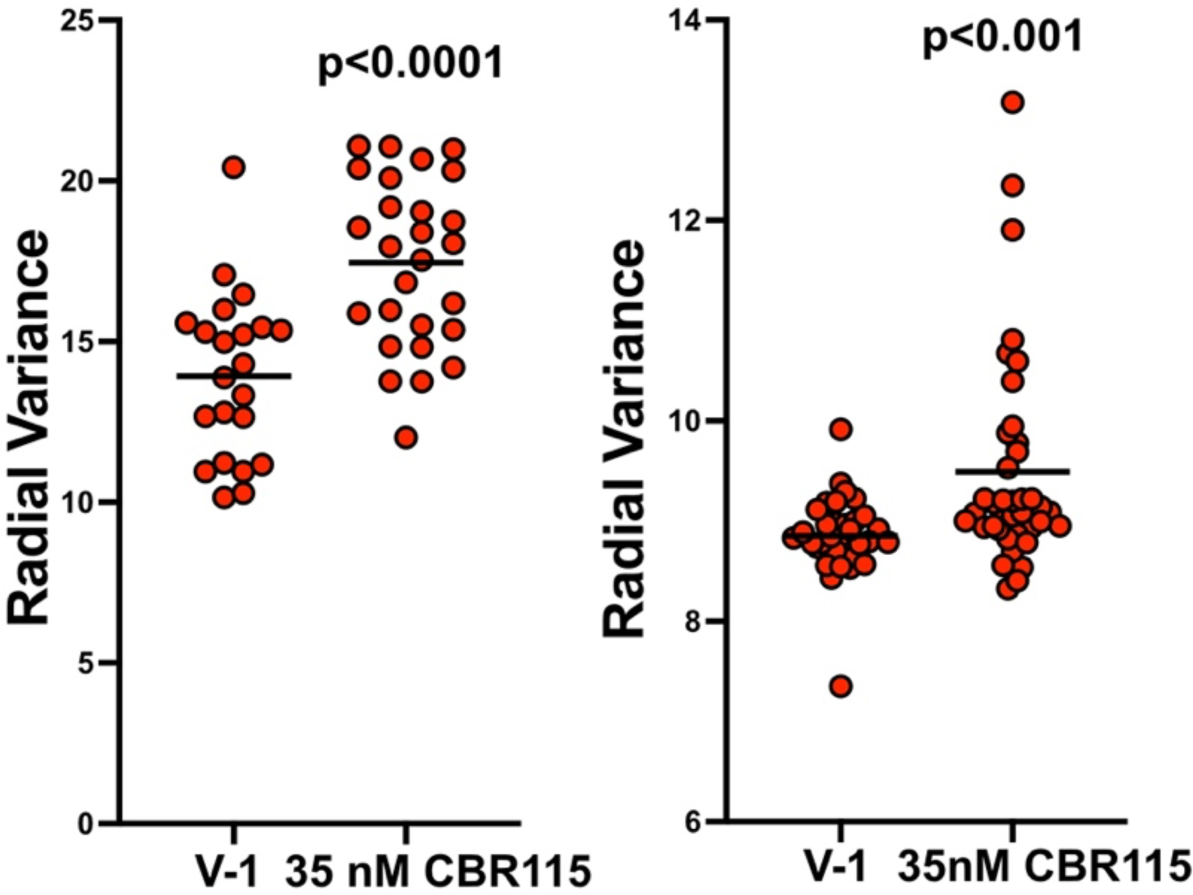
Radial variance increases with CBR115 CARMIL. V-1 alone is compared with CBR115 CARMIL plus V-1. Radial distances from the center of each bead were determined and a spread of the distances gave the radial variance value for actin fluorescence. Two independent experiments are shown. Non-parametric t-test (Mann-Whitney) p values are shown.

**Supplementary Figure 3.**
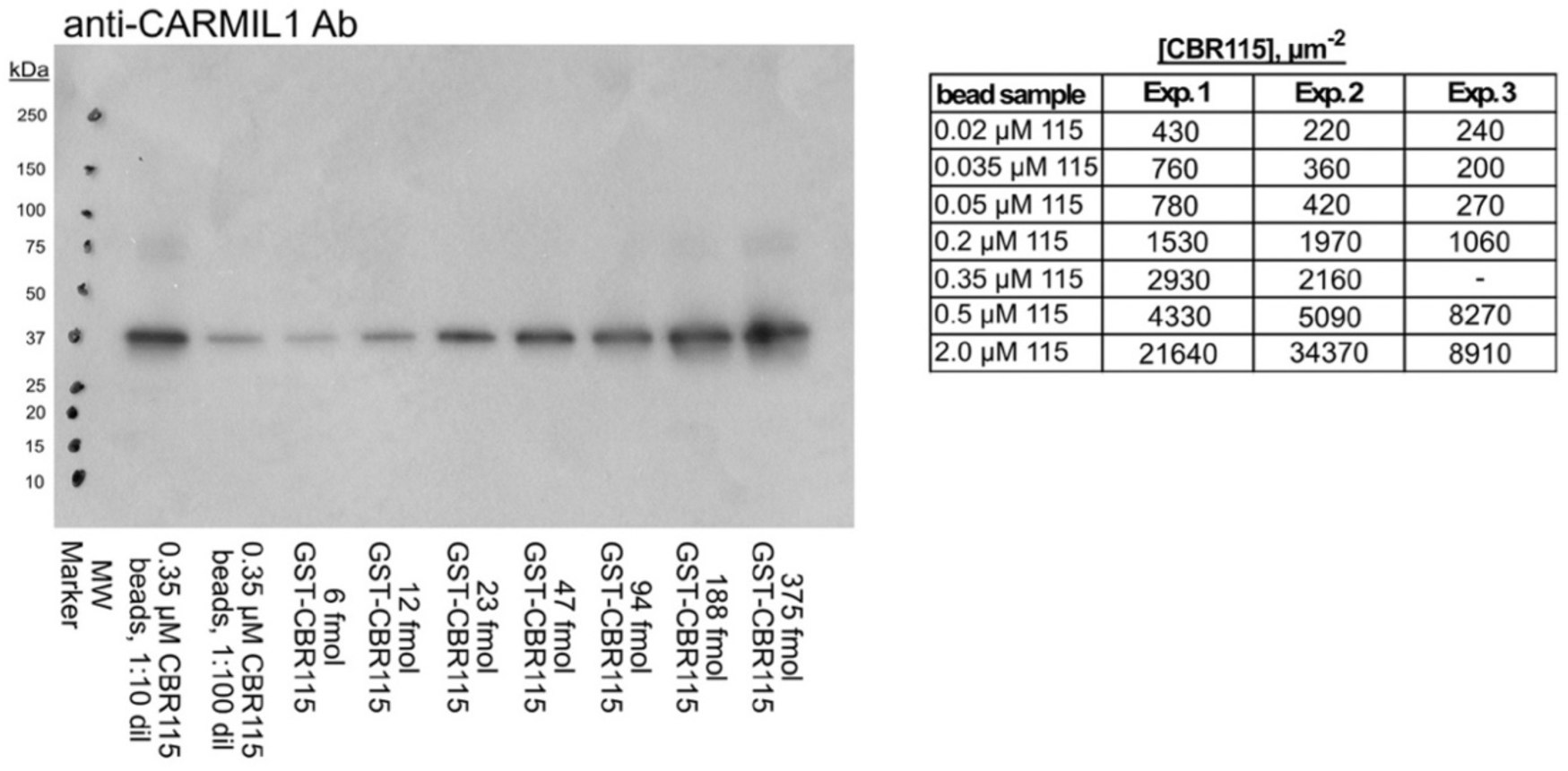
Western blot image for quantitation of GST-CBR115 on beads. One representative image. Positions of molecular weight standards are indicated on the left side. Bead samples at two different dilutions are in the left two lanes. Standards with known amounts of purified GST-CBR115 are in the seven lanes on the right, in amounts as indicated. Table lists amounts of CBR115 per µm^2^ of bead surface for three values from three blots.

**Supplementary Figure 4.**
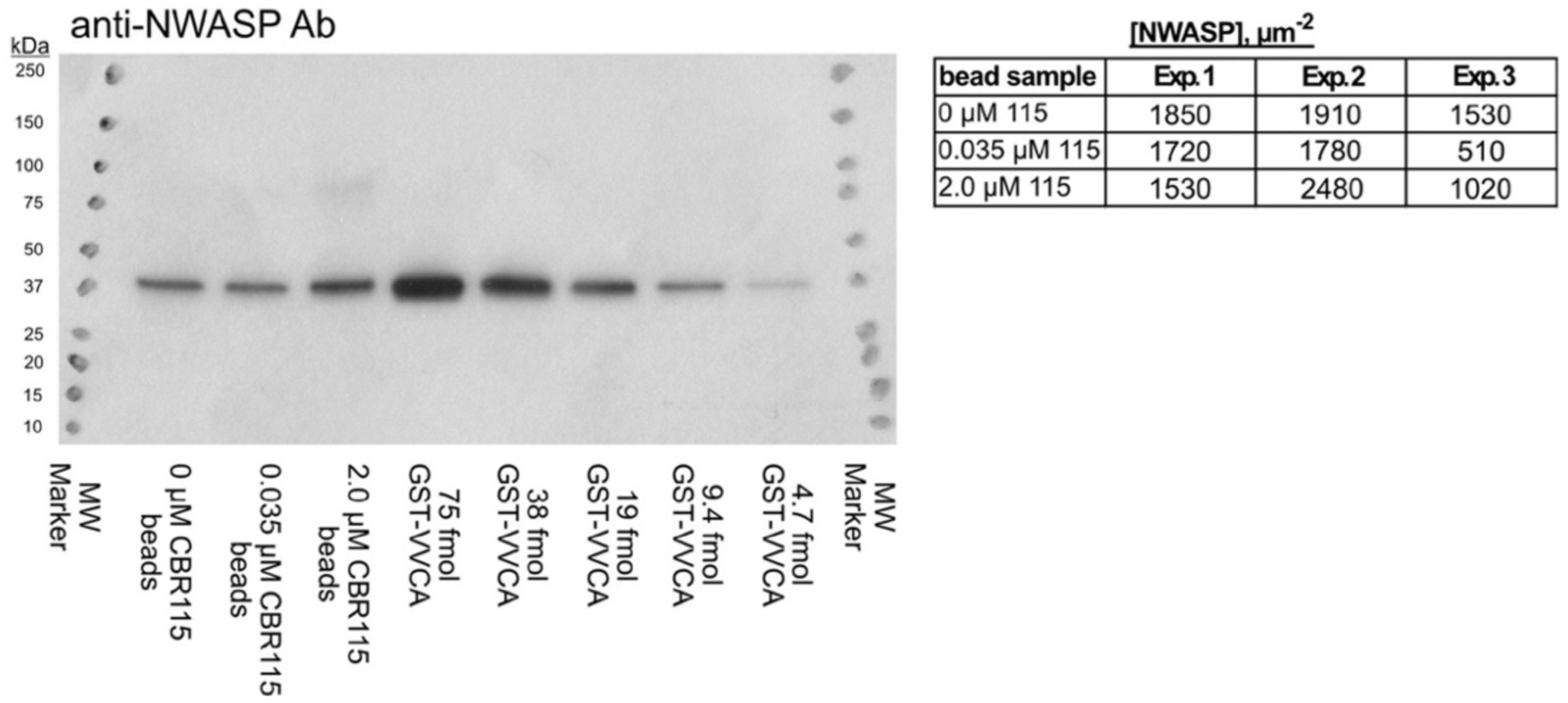
Western blot image for quantitation of GST-VVCA on beads. One representative image. Positions of molecular weight standards are indicated on both sides. Bead samples from three different bead samples are in the left three lanes. Standards with known amounts of purified GST-VVCA are in the five lanes on the right, in amounts as indicated. Table lists amounts of GST-VVCA per µm^2^ of bead surface for three values from three blots.

**Supplementary Figure 5.**
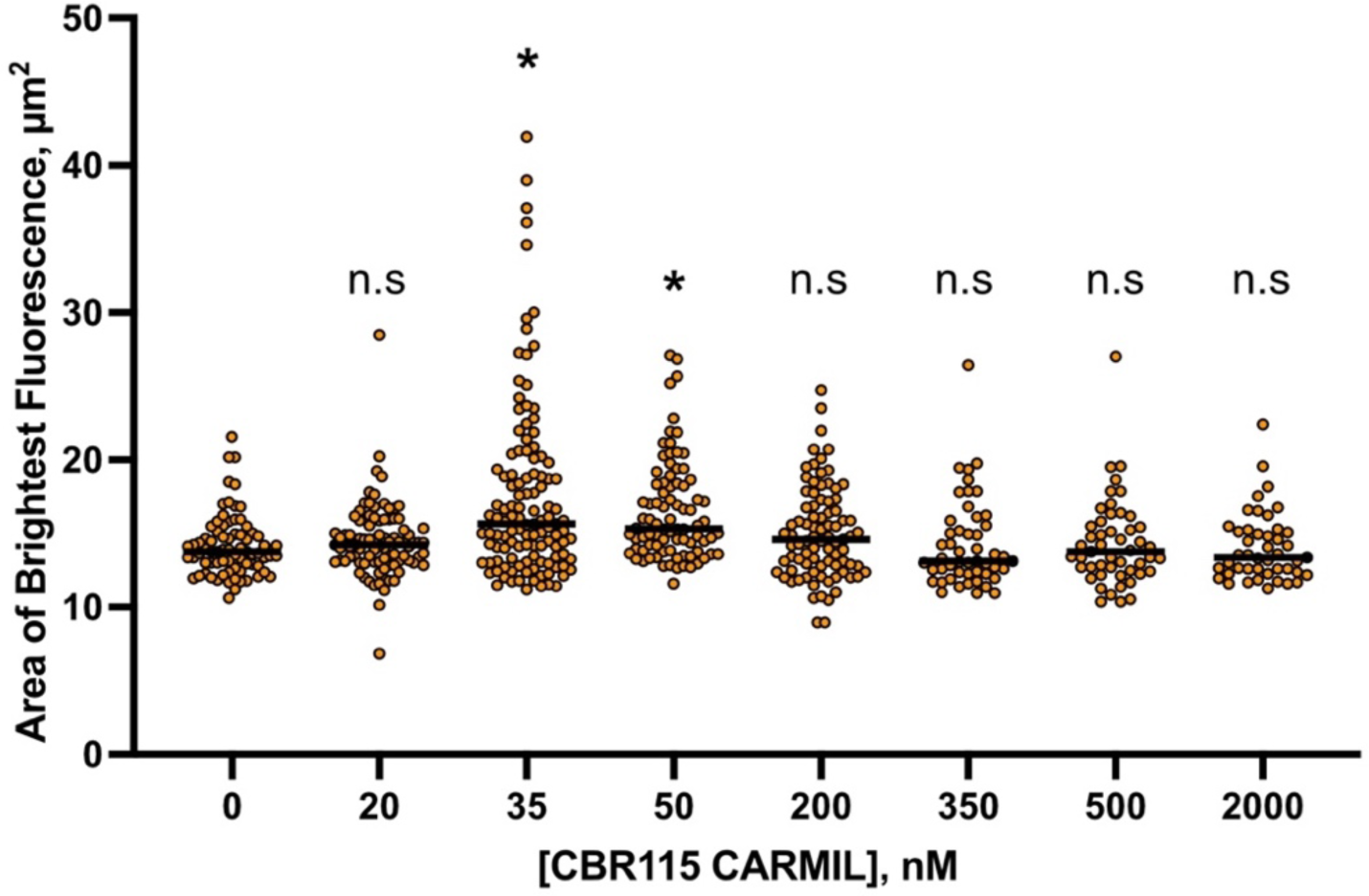
Area of brightest actin fluorescence surrounding beads increases with small amounts of CBR115 CARMIL. Thresholding was set so that only the brightest actin fluorescence was included, excluding the less bright actin clouds. Data are from three independent experiments. Non-parametric t-test (Mann-Whitney) p values for data compared to 0 nM CBR115 conditions are indicated: * is p<0.0001, n.s. is non-significant.

**Supplementary Figure 6.**
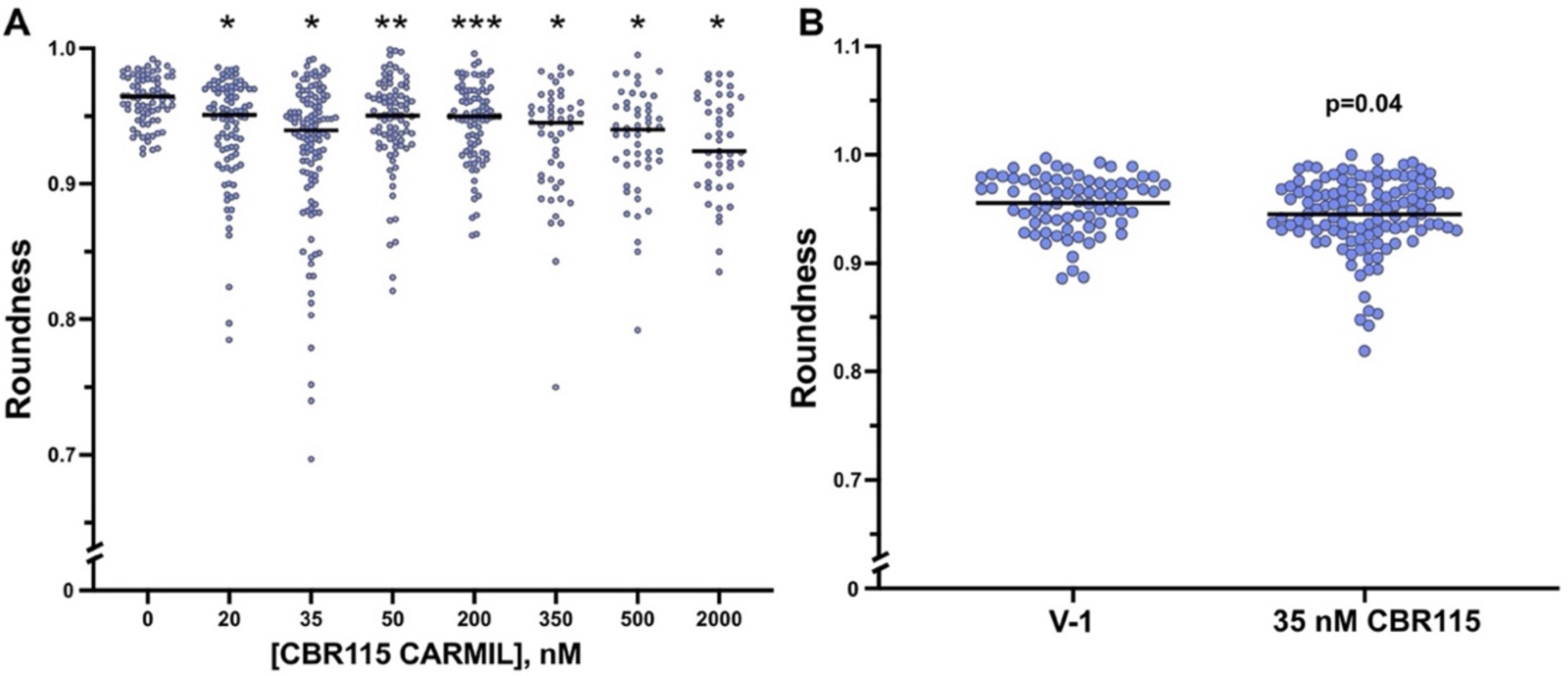
CBR115 CARMIL alters the shape of the actin closest to the bead surface. **A.** Roundness of the brightest actin fluorescence closest to the bead surface is plotted for each GST-CBR115 concentration. Data are from three independent experiments. Non-parametric t-test (Mann-Whitney) p values with data compared to 0 nM CBR115 conditions are indicated: * is p<0.0001, ** is p<0.001, ** is p=0.0001. **B.** Roundness of the total actin fluorescence surrounding the beads including the diffuse actin cloud. Data are from three independent experiments. Non-parametric t-test (Mann-Whitney) p value shown.

**Supplementary Figure 7.**
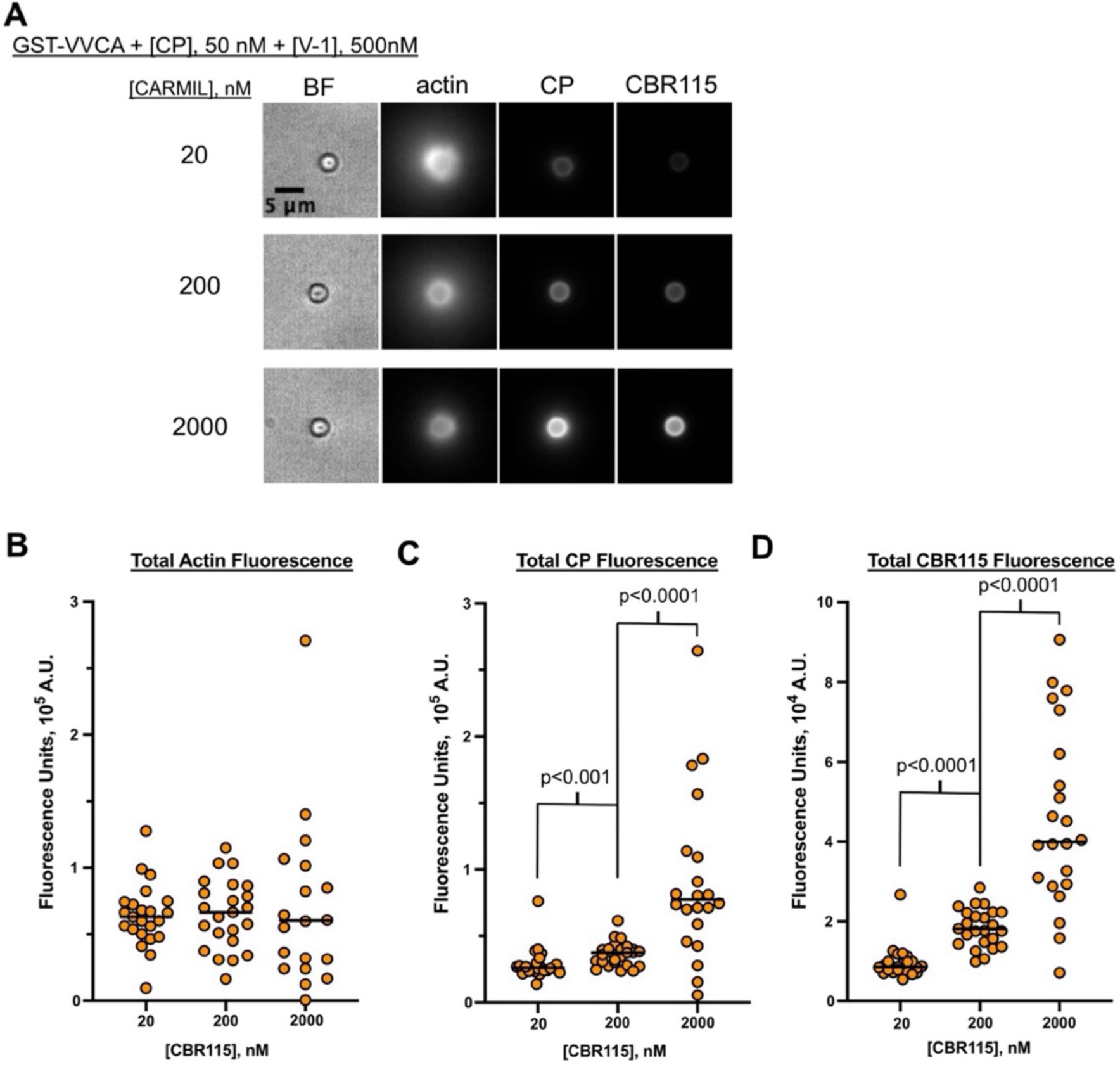
CBR115 CARMIL recruits CP to beads and depends on concentration. **A.** Beads were prepared and incubated with reaction mixture, including Alexa 488 actin and Alexa 594 CP. At 30 min, reactions were arrested with phalloidin and Latrunculin B. **B and C.** Total fluorescence was measured as described in Methods. Non-parametric t-test (Mann-Whitney) p values are shown. **D.** To visualize the amount of GST-CBR115, beads were labeled with an anti-CARMIL1 Ab. Data shown are from two independent experiments. Non-parametric t-test (Mann-Whitney) p values are shown.

## Methods

### Buffers

#### Assay Buffer

20 mM HEPES, 100 mM KCl, 1 mM MgCl_2_, 1 mM EGTA, 1 mM TCEP, 1 mM ATP (pH 7.0). For bead actin polymerization microscopy assays, 0.2% methylcellulose (cP 400), 2.5 mg/ml BSA, and 20 mM beta-mercaptoethanol were added to help reduce Brownian motion, non-specific sticking to the coverslip surface, and photobleaching, respectively.

### Proteins

Porcine brain Arp2/3 complex was purchased from Cytoskeleton (Cat. No. RP01P, Denver, CO) and reconstituted per company’s instructions. Reconstituted protein was used within a month; otherwise, activity was diminished. Human profilin-1 (UniProt P07737) was a gift from Dr. Silvia Jansen (Dept. of Cell Biology & Physiology, Washington University, St. Louis) and was purified using previously published methods ^61^.

Rabbit skeletal muscle α-actin (UniProt P68135, *Oryctolagus cuniculus*) was extracted from muscle acetone powder (Pel-Freez) with 2 mM Tris-HCl, 0.2 mM CaCl2, 0.2 mM ATP, 0.5 mM DTT, 1 mM NaN3 (pH 8.0) (G-Ca buffer), and then polymerized by addition of 50 mM KCl and 2 mM MgCl_2_. Tropomyosin was removed by further incubation in the presence of 0.8 M KCl. The F-actin was pelleted by centrifugation and was depolymerized by dialysis against G-Ca buffer. The G-actin was gel-filtration purified on Sephacryl S-300 in G-Ca buffer. Purified G-actin was dialyzed into 25 mM HEPES, 0.1 M KCl, 2 mM MgCl_2_, 1 mM NaN_3_, 0.2 mM ATP (pH 7.0), then 0.8 mol of N-(1-pyrene) maleimide or 1.0 mol of Alexafluor-488-maleimide was added per mol actin, and the solution was incubated overnight at 4°C. The labeled actin was pelleted by centrifugation, and the pellets were dissolved and depolymerized by dialysis against G-Ca buffer.

Mouse CPα1β2 heterodimer (UniProt Q5RKN9 and Q923G3, pBJ 2041), assembled from the two co-expressed subunits, was purified as described ^41^. Human V-1 (UniProt P58546, pBJ 2438) was purified as described ^41^.

N-terminal GST-tagged human CARMIL1 (UniProt Q5VZK9) fragment E964 to S1078 (aka “CBR115”, pBJ 2411) was purified as described ^62^. N-terminal GST-tagged human N-WASp fragment P392 to D505 (pBJ 2497, WH2-WH2-CA, aka “VVCA”) was composed of (GST-GSGSGS(linker) – LEVLFQGP (PreScission protease site) – N-WASp (UniProt O00401) P392 to D505). Codon-optimized DNA was synthesized and inserted into a pRSF-1b vector (by Azenta Life Sciences, Burlington, MA). The GST fusion protein was expressed in E. coli BL21 Star (DE3) and affinity-purified on a Glutathione Sepharose 4 Fast Flow column (GE Healthcare), followed by ion-exchange chromatography on a POROS GoPure HQ column (Applied Biosystems) in 20 mM Tris-HCl, 1 mM NaN3, 10 mM DTT (pH 7.5) with elution by a linear gradient to 1 M NaCl. Fractions containing the purified GST fusion protein were pooled, concentrated, and stored at −70 ^°^C.

For experiments imaging the location of CP around the bead, we used fluorescently tagged CP, prepared as follows. In a previous study, CP was mutated to change all Cys residues to Ser (Cys-null CP), and Cys-null CP was shown to retain biochemical activity ^30^. We introduced Cys residues at two specific sites by changing Ser residues – residue 9 of CPα1 and residue 161 of CPβ2 (Cys-null CP with α1 S9C β2 S161C) ^30^. We then prepared and biochemically characterized double-Alexa 594-labeled CP α1 S9C β2 S161C. Double-Alexa 594-labeled Cys-null CP with α1 S9C β2 S161C (pBJ 2478) was prepared as described ^30^. Fluorescent CP was used only for localization experiments, not for actin assembly reconstitution experiments.

### Protein-Coating of Beads

Glutathione-functionalized beads with a diameter of 2 µm (Spherotech, GSHP-20-5, Lake Forest, IL) were sonicated for 10 min, centrifuged (10 min, 5000 rpm), and washed using assay buffer (no ATP). Beads were suspended in assay buffer (no ATP) containing the following concentrations of GST fusion proteins: 2 µM GST-VVCA, 0-2 µM GST-CBR115, 0-2 µM GST. GST alone was used as a placeholder when the concentration of GST-CBR115 was less than 2 µM, so that the total concentration of GST was 4 µM in all samples during the coating process. Beads were incubated with GST fusion proteins for 3 hours on ice, with frequent agitation to maintain the beads in suspension. Beads were centrifuged and washed once in assay buffer (no ATP) with 1% BSA, followed by a second wash step in assay buffer (no ATP) with 0.1% BSA. Protein-coated beads were then suspended in assay buffer with 0.1% BSA and 1 mM ATP. Beads were counted, and a standard curve of OD600 values for bead suspensions of known concentrations was generated. An OD600 reading was taken for each protein-coupled bead sample to quantify the amount of beads, and the volume was adjusted for each sample so that the same number of beads could be used for each reaction (∼3.2×10^5^ beads / µL stock, with 1 µL per reaction). Protein-coated beads were used within one or two days.

We determined the amount of protein coupled to the beads. Bead suspensions were passed through a 0.2 µm centrifugal filter (ThermoFisher, 50-197-9571). The beads were collected without washing, and SDS-PAGE sample buffer was added. These samples of bead-bound protein were electrophoresed on an SDS gel, transferred to PVDF membrane, and probed with a 1:1000 dilution of anti-CARMIL1 polyclonal antibody (ThermoFisher, 27133-1-AP) or anti-N-WASP polyclonal antibody (ThermoFisher, PA5-52198). The secondary antibody was a 1:10,000 dilution of peroxidase-conjugated goat anti-rabbit IgG (Sigma, A6154). Blots were developed with chemiluminescence substrate (Thermo Scientific # 32209). Autoradiograms were collected and digitized, and intensities of bands were analyzed with the gel analysis tool of ImageJ ^63^. Standard curves were generated using known amounts of GST-CBR115 or GST-VVCA electrophoresed on the same gel with bead samples.

### Bead actin polymerization microscopy assays

For bead-based actin polymerization assays, protein-coated beads (∼3.2×10^5^ beads or ∼1.6×10^4^ beads / µL) were mixed with 100 nM Arp2/3 complex, CP (0-200 nM), 5 µM profilin, 5 µM actin (10% Alexa 488-labeled actin), and V-1 (0-5000 nM) in assay buffer containing 0.2% methylcellulose (cP 400), 2.5 mg/ml BSA, and 20 mM β-mercaptoethanol. To ensure that relevant complexes formed before the start of the actin polymerization reaction, we pre-mixed three pairs of components: Arp2/3 complex with beads, actin monomers with profilin, and CP with V-1. The pairs were incubated for 15 min at RT. All three of the pre-mixes were then combined into one vessel to start the polymerization reaction. After 15 min at RT, the reaction mixture was mounted between a coverslip and a glass slide, sealed using clear fingernail polish and imaged after another 15 min. Alternatively, the reaction was arrested after 30 min of incubation by adding 10 µM phalloidin and 10 µM Latrunculin B, mounted between a coverslip and a glass slide, and imaged immediately. These reaction times were chosen after pilot experiments testing a range of times.

Samples were imaged using a 60x 1.45 NA oil objective on an inverted Olympus IX81 microscope using a mercury lamp for excitation. Images were collected in wide-field fluorescence and bright-field modes with a Hamamatsu EM-CCD (C9100) camera using Micro-Manager software ^64^. Digital images were analyzed using Fiji software ^65^.

### Antibody labeling of bead samples for imaging

Bead samples were prepared as described above, and reactions were arrested at 30-min with 10 µM phalloidin and 10 µM Latrunculin B (arrest mixture in assay buffer, no ATP). Beads were pelleted by centrifugation (10 min, 5000 rpm), suspended in arrest-mixture assay buffer (no ATP) with 3% BSA and 1:400 anti-CARMIL1 antibody (ThermoFisher, 27133-1-AP). After incubation for 2 hr at RT, beads were centrifuged and washed twice using arrest-mixture assay buffer (no ATP). Secondary antibody (1:400 Alexa 647-labeled goat anti-rabbit, ThermoFisher A21244) was added in arrest-mixture assay buffer (no ATP) with 3% BSA. After incubation beads for 1 hour at RT, beads were washed twice, resuspended, and imaged as above.

### Pyrene actin polymerization assays

Pyrene actin polymerization with CP titration (Figure 1, panel D) was performed in assay buffer in 0.2 mL mixture of the following in assay buffer: actin monomers (5 µM, 5% pyrene-labeled), profilin (5 µM), GST-VVCA beads (1.2×10^4^ beads/µL), and Arp2/3 complex (100 nM). CP was present at concentrations from 0 to 200 nM. GST-VVCA beads, Arp2/3 complex, and CP were pre-mixed for 20-60 min at RT. Pyrene-labeled actin and profilin were pre-mixed for ∼10 min at RT and then added to the bead mixture at time zero. For the V-1 titration (Figure 2, panel D), conditions were similar except as follows: GST-VVCA beads, Arp2/3 complex, CP and V-1 were pre-mixed. Final concentrations were GST-VVCA beads at 3×10^4^ beads/µL, CP at 0 or 50 nM, and V-1 at concentrations from 0 to 20 µM. Pyrene fluorescence was measured using time-based scans on a plate reader (Biotek Synergy H4 Hybrid Multi-Mode Microplate Reader with Gen5 software, BioTek Instruments, Winooski, VT) at 25°C with excitation at 365 nm and emission at 407 nm.

### Analysis of fluorescence images of beads

To measure the area of actin fluorescence, two methods were employed. When beads displayed asymmetric tail growth, the area of fluorescence was easily identified by intensity, and an observer outlined the area using the freehand drawing tool and the Measure tool in Fiji. When beads were surrounded by diffuse clouds of actin fluorescence, the edge of the actin cloud was less clear, so a threshold level was set and used as a mask to demarcate the edge. The freehand drawing tool was used to outline around the threshold mask to determine the area of the actin cloud.

To measure the total actin fluorescence associated with each bead, a 20×20 µm box was drawn centered around the bead, and the value of the integrated density was recorded. Next, a 30×30 µm box was drawn around the outside of the 20×20 µm box to determine the background value surrounding each bead. The background was calculated using the following equation:

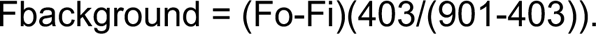

In this equation, Fo is the outer box fluorescence and Fi is the inner box fluorescence. The total fluorescence of the bead was then calculated as:

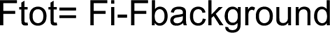

To determine the shape of the fluorescence, we used thresholding to determine the area of brightest fluorescence surrounding the beads, or total fluorescence to include the clouds. Once the threshold was set, we used the Analyze Particles tool in ImageJ to determine the aspect ratio and roundness of each shape. Roundness is mathematically defined as the inverse of the aspect ratio.

#### Image Acquisition and Preprocessing

TIFF images of beads were cropped manually to isolate beads in focus. The cropped images were processed using Python, leveraging libraries such as PIL, NumPy, and skimage. Each image was converted to a grayscale array and subjected to an adaptive thresholding algorithm to separate the foreground (bead with actin cloud) from the background. The threshold was dynamically calculated based on the mean and standard deviation of pixel intensities, where pixels greater than the mean plus two standard deviations were considered part of the foreground.

#### Object Segmentation and Feature Extraction

Connected components in the binary images were identified using the label function from skimage. Properties of these labeled regions, such as area and perimeter, were extracted using regionprops. The largest connected component was isolated for further analysis to focus on the bead in each image.

#### Morphological Analysis

For the largest object in each image, morphological features, including aspect ratio and roundness, were calculated. Aspect ratio was defined as the ratio of the major to minor axis lengths of the object. Roundness is the inverse of the aspect ratio. The centroid of the object was computed to facilitate distance measurements from this central point to all other points within the object.

#### Radial Variance Analysis

A grid of coordinates was generated for the image, and distances from each point to the centroid were calculated. Distances within the object were isolated using a masked pixel array, and the variance of these distances (radial variance) was computed to quantify the dispersion of the object’s mass around its centroid.

Let (x_i_, y_i_) be the coordinates of the i-th point within a region of interest, and (x_c_, y_c_) be the coordinates of the centroid of this region. The radial distance d_i_ from each point to the centroid is calculated using the Euclidean distance formula:

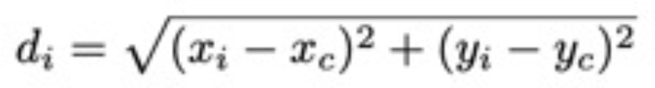

For a set of n points, the radial variance σ^2^ is computed as the variance of these radial distances:

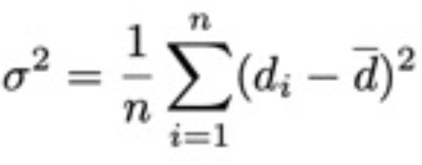

where *d̄* is the mean radial distance, calculated as :

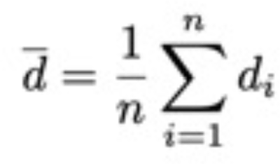

#### Visualization

Visual representations included plotting the binary mask of the largest object with the centroid marked, displaying a histogram of radial distances within the object, and creating a polar plot of the histogram to visually assess the distribution of pixel distances from the centroid.

#### Radial Intensity Scan

The original grayscale image was used to assess the intensity distribution relative to the centroid. Intensity values at each pixel within the largest object were mapped against their respective distances from the centroid. These values were binned, and the mean intensity for each bin was calculated along with its standard deviation. A plot of radial intensity versus distance provided insight into the intensity profile of the actin cloud.

#### Variance Metric

A novel variance metric was calculated by dividing the sum of standard deviations of intensity values by the sum of mean intensities across all bins. This metric aimed to quantify the intensity variability relative to the average intensity, providing a measure of heterogeneity within the object. Each of these steps was iterated for all TIFF images in the directory, compiling an array of metrics that summarized the morphological features.

## Abbreviations

CP: actin capping protein.
F-actin: filamentous actin.
CPI: capping-protein-interacting.
CARMIL: capping protein, Arp2/3, and myosin I linker.
NPF: Nucleation promoting factor.

## Notes

### Competing Interest Statement

The authors have declared no competing interest.

